# Forage conservation is a neglected nitrous oxide source

**DOI:** 10.1101/2024.03.19.585631

**Authors:** Seongmin Yang, Maheen Mahmood, Rudra Baral, Hui Wu, Marc Almloff, Lauren E. Stanton, Doohong Min, Brenda K. Smiley, J. Chris Iiams, Jisang Yu, Jeongdae Im

## Abstract

Agricultural activities are the major anthropogenic source of nitrous oxide (N_2_O), an important greenhouse gas and ozone-depleting substance. However, the role of forage conservation as a potential source of N_2_O has rarely been studied. We investigated N_2_O production from the simulated silage of the three major crops—maize, alfalfa, and sorghum—used for silage in the US, which comprises over 90% of the total silage production. Our findings revealed a substantial N_2_O could be generated, which potentially places forage conservation as the third largest N_2_O source in the agricultural sector. Notably, the application of chlorate as an additive significantly reduced N_2_O production, but neither acetylene nor intermittent exposure to oxygen showed any impact. Overall, the results highlight that denitrifiers, rather than nitrifiers, are responsible for N_2_O production from silage, which was confirmed by molecular analyses. Our study reveals a previously unexplored source of N_2_O and provides a crucial mechanistic understanding for effective mitigation strategies.

**Significance Statement:** N_2_O is the third most important greenhouse gas (GHG) and agriculture contributes 80% of the total anthropogenic emissions in the US. The major sources of N_2_O in the agricultural sector identified by the USEPA include agricultural land management, manure management, and the field burning of agricultural residues. Here, we show that forage conservation could be a significant unaccounted source of N_2_O, surpassing the field burning by 30. Our study provides a mechanistic understanding of N_2_O production and a simple and effective remedy for reducing N_2_O emissions. The findings have substantial implications for mitigating climate change, informing policy-makers, and guiding future research on reducing greenhouse gas emissions from livestock production.

---

Nitrous oxide (N_2_O) is the third most important greenhouse gas (GHG), following carbon dioxide (CO_2_) and methane. On a molar basis, N_2_O has a warming potential 300 times greater than CO_2_ and remains in the atmosphere for an extended period, estimated at 100–150 years (1). Recent studies have also emphasized N_2_O as the primary ozone-depleting substance in the stratosphere (2, 3). Atmospheric N_2_O concentrations have increased by over 20% compared to preindustrial levels, with the fastest increase occurring in the last five decades (4, 5). While natural processes and human activities contribute to N_2_O production, agricultural activities are one of the dominant sources, responsible for over two-thirds of global anthropogenic N_2_O emissions (5–7). Numerous countries compile GHG emission inventories following the technical guidelines provided by the Intergovernmental Panel on Climate Change. For example, the United States Environmental Protection Agency (US EPA) annually publishes a report titled “Inventory of U.S. Greenhouse Gas Emissions and Sinks,” which accounts for N_2_O emissions from agricultural activities through three sources: agricultural land management, manure management, and the burning of agricultural residues.

Forages, the plant materials consumed by herbivores, are conserved to sustain livestock during periods of limited pasture growth or inadequate grazing conditions (8, 9). Globally significant for productive and efficient livestock production, forage conservation methods mainly involve hay and silage. For long-term storage, hay is dried to below 20% moisture (12%–20%, w/w) to curtail microbial growth and stored under aerobic conditions (8). Conversely, silage is produced at higher moisture levels (40%–70%, w/w) and stored strictly under anaerobic conditions to facilitate fermentation. Indigenous or exogenous lactic acid bacteria (LAB) convert soluble carbohydrates into organic acids, predominantly lactic acid, which acts as a natural preservative that inhibits unwanted microorganisms (10, 11). Commercial silage inoculants containing LAB, such as *Lentilactobacillus buchneri* and other facultative anaerobic bacteria, are used as additives to enhance silage fermentation (12). The market for global silage inoculants reached USD 503 million in 2021 and is projected to grow to USD 630 million by 2028 at a compound annual growth rate of 3.8% (13). Silage is a significant segment of the global livestock industry, with 162.3 million metric tons (MMT) harvested in the US in 2022 (14, 15).

Despite its high production volume and extensive use, forage conservation has been limitedly studied as a potential source of GHG emissions (15). While gas production during forage conservation has been investigated, previous studies have primarily focused on odorous chemicals, such as volatile organic compounds and ammonia (NH_3_) (16–18). At best, the detection of N_2_O has been reported in previous studies (19–22), but a quantitative assessment is still lacking. That is, our study, to our knowledge, is the first to provide comprehensive quantitative estimates on a per-crop basis, which can be scaled to national- and sector-level estimates for N_2_O emission from forage conservation.

The precise mechanism underlying N_2_O production remains partially understood, but microbial processes are widely considered the main contributors to N_2_O emissions (5, 23–26). While nearly all microorganisms involved in the biogeochemical nitrogen cycle can potentially produce N_2_O, specific microbial pathways including heterotrophic denitrification, NH_3_ oxidation, and nitrifier denitrification, are pivotal to N_2_O production (26, 27). Heterotrophic denitrification is a multistep respiration process that involves the reduction of oxidized mineral forms of nitrogen (i.e., nitrate (NO_3_^-^) and nitrite (NO_2_^-^)) to gaseous nitric oxide (NO), N_2_O, and dinitrogen (N_2_). This process typically occurs under anaerobic conditions, although recent discoveries have identified a new group of aerobic denitrifying bacteria (28). Conversely, NH_3_ oxidation and nitrifier denitrification occur under aerobic conditions. Nitrification involves a two-step process: the oxidation of NH_3_ to NO_2_^-^ by ammonia-oxidizing bacteria (AOB) and archaea (AOA), followed by further oxidation to NO_3_^-^ by NO_2_^-^-oxidizing bacteria. N_2_O is indirectly produced through the chemical decomposition of intermediate or end products of NH_3_ oxidation (hydroxylamine, nitroxyl hydride, or NO_2_^-^) (29). Additionally, certain AOB can directly produce N_2_O through nitrifier denitrification by oxidizing NH_3_ to NO_2_^-^ and subsequently reducing it to NO and N_2_O (30–32).

In this study, we conducted a comprehensive quantitative assessment of N_2_O emissions during forage conservation, especially in silage form. Simulated silages derived from three major silage crops in the US—maize, alfalfa, and sorghum—were monitored for N_2_O emissions over a 4-week period. Our findings showed significant N_2_O release from silages, making forage conservation the third-largest source of N_2_O emissions within the agricultural sector. Further experiments confirmed the significant role of denitrification in N_2_O production in conserved forages, as validated by molecular analyses.

## Results

### Chemical properties of silage materials

The freshly chopped, non-inoculated plant materials showed characteristic nutritional properties for each crop (SI Fig. S1). Among them, alfalfa, a leguminous crop, showed notably higher concentrations of proteins and amino acids than maize and sorghum. Conversely, cereal crops, such as maize and sorghum, exhibited a higher starch content than alfalfa. The total protein and amino acid contents in alfalfa decreased as it matured, accompanied by an increase in fiber (acid detergent fiber (ADF) and neutral detergent fiber (aNDF)) and lignin contents. Notably, the alfalfa variety HybriForce 3400 consistently showed higher protein and amino acid contents than HVX MegaTron, regardless of maturity stage.

### N_2_O emissions from simulated silage

The total amount of N_2_O produced in the maize, alfalfa (HybriForce 3400, harvested at mid-bud stage, Cv_2_ Hv_1_), and sorghum silage over 28 d of incubation was 6.7 (*±*0.7), 62.3 (*±*4.0), and 1.8 (*±*0.1) mL, which corresponded to 18.2 (*±*1.9), 169.7 (*±*10.9), and 4.8 (*±*0.2) g CO_2_ equivalent (eq.) per kg_DM_, respectively (Fig. 1a). N_2_O emissions began immediately after incubation commenced, with the majority (> 90%) being produced within 24 h for maize and sorghum and 5 d for alfalfa. For alfalfa harvested at the same growth stages, no significant difference in N_2_O production was observed between the varieties (*p* > 0.05) (Fig. 1b). However, incubations with alfalfa harvested at the later maturity stages (i.e., early flowering stage, Hv_2_) produced significantly lower N_2_O emissions (*p* < 0.05) regardless of the varieties (Fig. 1b). The statistical analysis is summarized in SI Fig. S2.

**Fig. 1.**
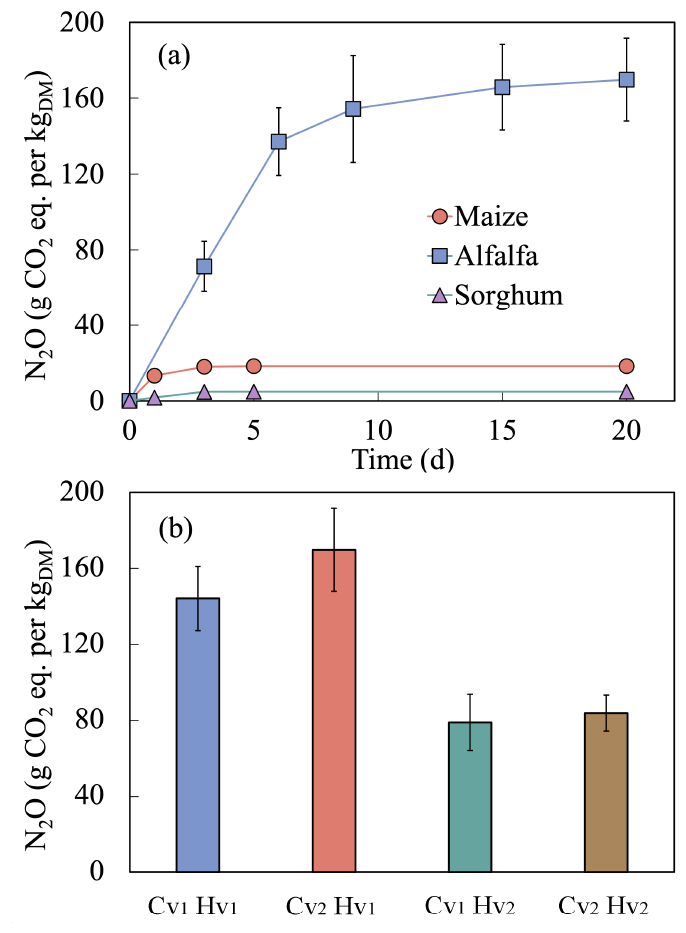
Cumulative N_2_ O production of (a) maize, alfalfa, and sorghum and (b) two distinct alfalfa varieties harvested at two different stages of maturity. The labels Cv_1_ and Cv_2_ denote the two alfalfa varieties, HVX MegaTron and HybriForce 3400, respectively. Similarly, the Hv_1_ and Hv_2_ denote alfalfa samples harvested at the mid-bud and early flowering stages, respectively. The error bars represent the standard deviations derived from the triple incubations. Error bars may not be visible if their magnitude is smaller than the symbols.

### Effects of different treatments on N_2_O production

The effects of various treatments on N_2_O production were examined within the same sample group, revealing consistent trends regardless of the crops or alfalfa varieties harvested at different growth stages (Fig. 2a). Compared to the controls (I^-^), the addition of inoculants (I^+^) had no significant effect on N_2_O production (*p* > 0.05), except for sorghum, where a significant difference was observed (*p* < 0.05). Notably, applying chlorate treatment and the inoculant (I^+^ Ch^+^) significantly reduced N_2_O production by up to 99%. The addition of acetate as an external carbon source alongside the inoculant (I^+^ Ac^+^) resulted in a statistically insignificant but numerically lower N_2_O production than I^+^ (*p* > 0.05). However, the effect of acetate on N_2_O reduction was significant in alfalfa Cv_2_ Hv_1_ and sorghum (*p* < 0.05). Conversely, neither acetylene (C_2_H_2_) addition nor intermittent O_2_ exposure at different periods (Days 3, 5, and 10) impacted N_2_O production (Fig. 2b). Lower chlorate concentrations, as low as 0.01% (w/w), still achieved 92% N_2_O reduction (Fig. 2c). Residual chlorate could not be quantified due to technical limitations in ion chromatography. The statistical analysis is summarized in SI Table S1.

**Fig. 2.**
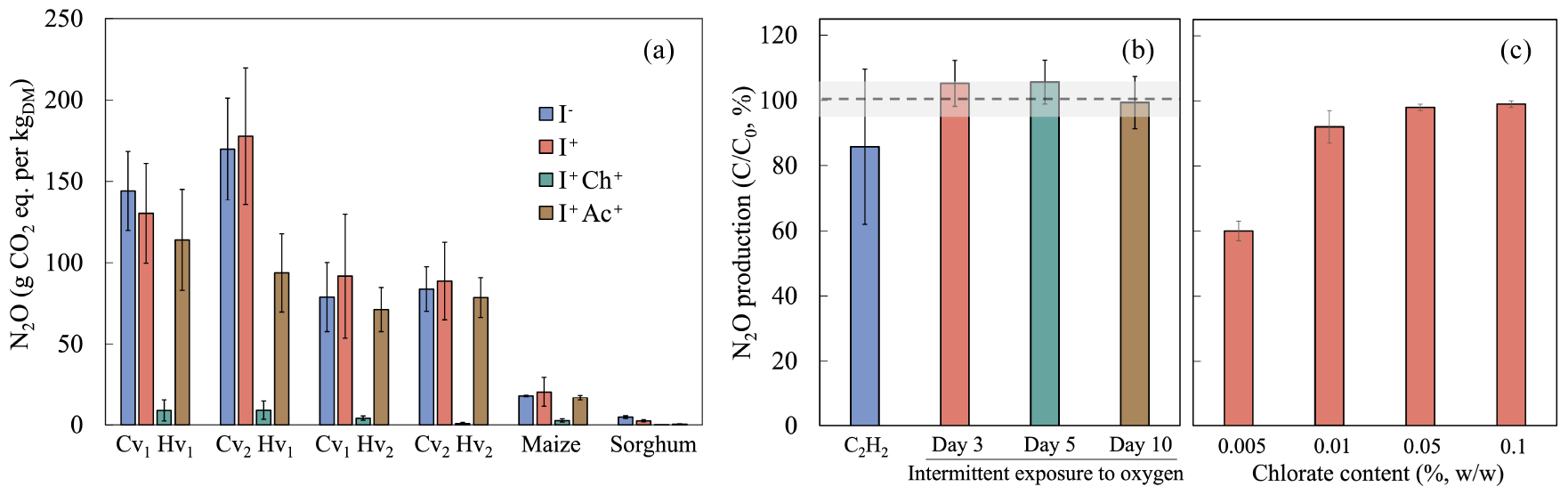
Effects of various treatments on N_2_ O production. (a) No inoculant (I^-^), crop-specific commercial silage inoculant (I^+^), inoculant and chlorate (I^+^ Ch^+^), and inoculant and acetate (I^+^ Ac^+^). (b) Acetylene (C_2_ H_2_) addition and intermittent exposure to oxygen at different periods. (c) Different chlorate concentrations. Oxygen was added to separate bottles on each injection date. The N_2_ O production was normalized by control, with the shaded area representing the standard deviation of the control. The labels Cv_1_ and Cv_2_ denote two alfalfa varieties, HVX MegaTron and HybriForce 3400, respectively. Similarly, the labels Hv_1_ and Hv_2_ denote alfalfa samples harvested at the mid-bud and early flowering stages, respectively. The error bars denote the standard deviations derived from the triple incubations.

### Analysis of the correlation between N_2_O production and fresh matter nutrient parameters

The relationship between various nutrient parameters and N_2_O production in the controls (I^-^) was measured using a Pearson correlation coefficient (Fig. 3). Parameters related to protein and amino acids, including crude protein (r = 0.98, *p* = 0.031), total amino acids (r = 0.99, *p* = 0.001), NO_3_^-^-N (r = 0.94, *p* = 0.017), NH_3_-N (r = 0.96, *p* = 0.010), and NDICP (r = 0.92, *p* = 0.029), exhibited strong correlations with N_2_O production (r > 0.8 and *p* < 0.05). Conversely, parameters related to carbohydrates and fats, including ADF, aNDF, lignin, starch, ethanol-soluble carbohydrates, and total fatty acids, exhibited no significant correlation with N_2_O production (r < 0.4 or *p* > 0.05).

**Fig. 3.**
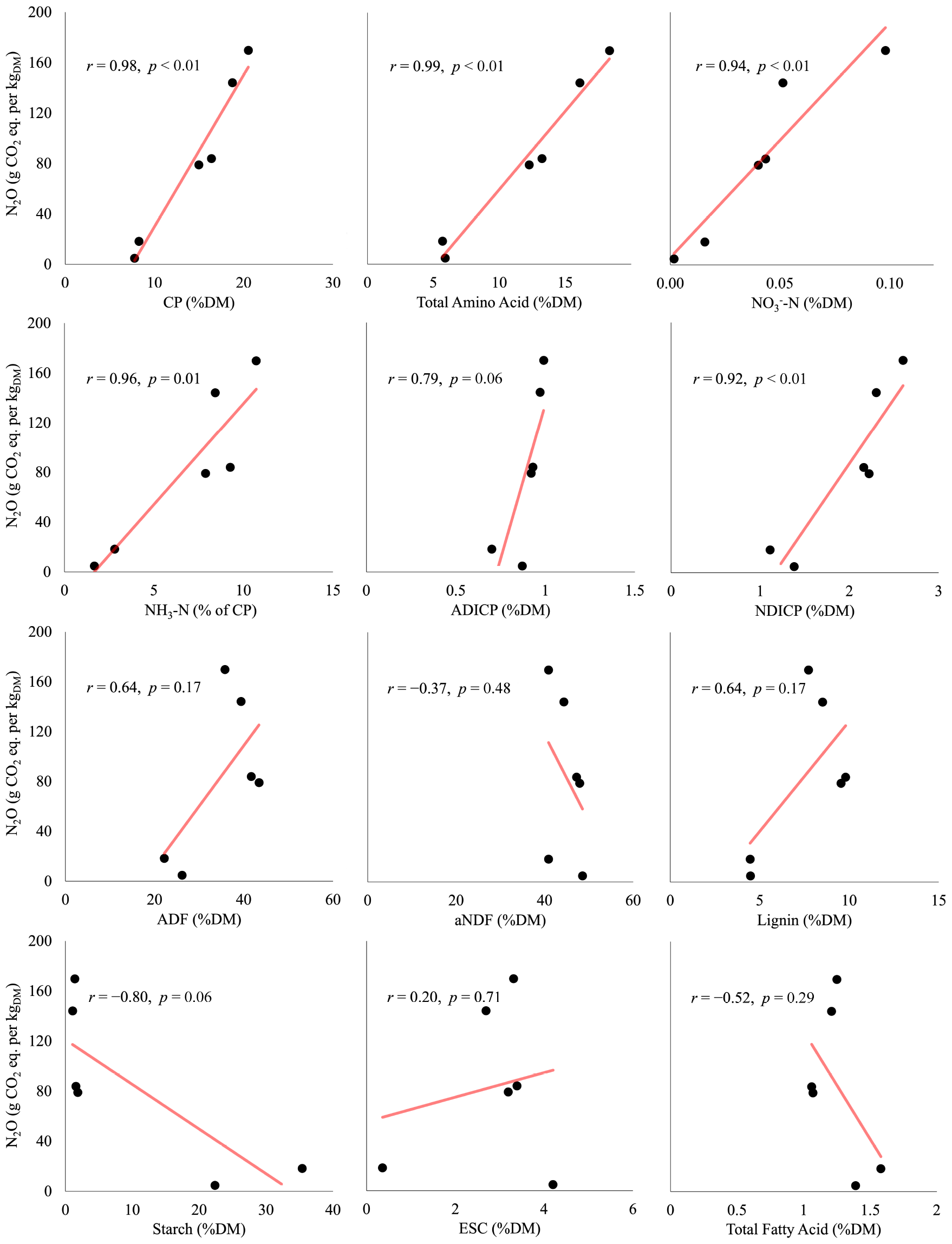
Correlation analysis of N_2_ O production and fresh matter nutrient parameters. DM: dry matter; CP: crude protein; ADICP: acid detergent insoluble CP; NDICP: neutral detergent insoluble CP; ADF: acid detergent treated fiber; aNDF: amylase treated neutral detergent fiber; ESC: ethanol soluble carbohydrates.

### Gene and transcript abundance dynamics

The abundance of the genes and transcripts in incubations with alfalfa harvested at the early flowering stage (Hv_2_) under various treatments is summarized in Fig. 4. Notably, the abundance of *narG*, the gene encoding membrane-bound nitrate reductase, exhibited a marked two-order-of-magnitude decrease with the addition of chlorate (I^+^ Ch^+^), in which N_2_O production was reduced by up to 99%, compared to I^+^. The abundance of other denitrification genes showed no trend over time across the various treatments. Notably, the abundances of bacterial and archaeal *amoA* gene encoding ammonia monooxygenase were lower than those of denitrification genes and no trend was observed over time across the various treatments. Transcript analysis revealed the expression of *narG* was completely suppressed by the addition of chlorate (I^+^ Ch^+^), similar to the gene abundance. Transcripts of archaeal and bacterial *amoA*, both *nir* genes encoding nitrite reductase, and *nosZ* gene encoding nitrous oxide reductase clade II were not detected. The expression level of *napA* gene encoding periplasmic nitrate reductase, *norB* gene encoding nitric oxide reductase, and *nosZ* gene encoding nitrous oxide reductase clade I were not affected by the treatments used.

**Fig. 4.**
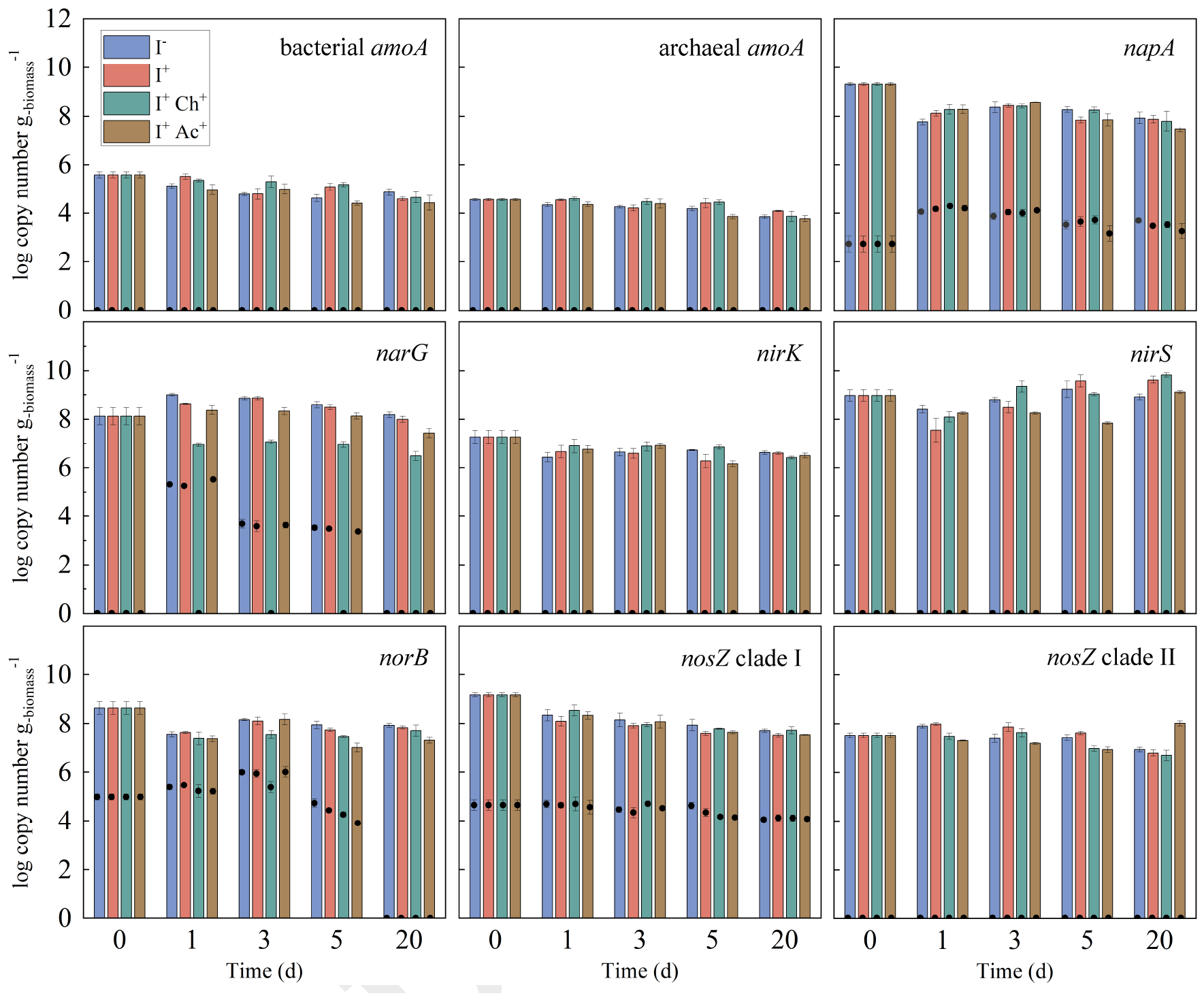
Abundance of functional genes (bar) and their transcripts (black circle) in incubation with alfalfa harvested at the early flowering stage (Hv_2_) under various treatments. The error bars represent the standard deviations of the triplicate incubation. Error bars may not be visible if their magnitude is smaller than the symbols.

## Discussion

### N_2_O production mechanism

The majority of N_2_O production in this study occurred within the first week of incubation (Fig. 1a), which is consistent with previous studies (21, 22). During this initial phase, transient aerobic or microaerobic conditions can be expected due to residual oxygen, followed by anaerobic conditions (33). Under these conditions, both ammonia oxidizers and denitrifiers are potential contributors to N_2_O production (34, 35). Ammonia oxidizers are recognized for producing N_2_O through direct (i.e., nitrifier denitrification) and indirect (i.e., abiotic processes) mechanisms. Furthermore, ammonia oxidizers can contribute to N_2_O production by converting NH_3_ to NO_3_^-^, subsequently fueling denitrification. However, N_2_O production did not increase when oxygen was added at different time points (*p* > 0.05, Fig. 3b), indicating that the contribution of nitrification to N_2_O production may not be significant. This finding is further supported by the observation that C_2_H_2_ (10 Pa), a potent inhibitor of bacterial and archaeal NH_3_ oxidation (36, 37), did not affect N_2_O production (*p* > 0.05, Fig. 3b). The absence of bacterial and archaeal *amoA* transcripts also confirmed the non-involvement of ammonia oxidizers in N_2_O production (Fig. 4). At high concentrations (1–20 kPa), C_2_H_2_ has been shown to inhibit N_2_O reductase activity of denitrifying microorganisms (38, 39). However, such inhibition (i.e., increased N_2_O production) was not observed in this study (Fig. 2b). The expression of Clade I *nosZ* remained unaffected in the samples supplemented with C_2_H_2_ (SI Fig. S3), suggesting that C_2_H_2_ did not inhibit denitrifying bacteria at the concentration used in this study (i.e., 10 Pa).

Due to the chemical similarities between NO_3_^-^ and chlorate, chlorate has been used as an inhibitor for dissimilatory NO_3_^-^ reduction, the first step in denitrification (40, 41). The significant decrease in N_2_O production, by up to 99% upon the addition of chlorate (Fig. 2a and SI Table S1), indicates that denitrifiers are the main contributors to N_2_O production. This finding was further confirmed by the diminished abundance of the *narG* genes and transcripts in the chlorate-amended samples (Fig. 4). In another study, the same concentration of chlorate (0.1% w/w) was used as a ruminant supplement to reduce *E. coli* O157:H7 population (42), suggesting that chlorate has the potential to be used as a silage additive to reduce N_2_O emissions. Further studies are warranted to assess the potential hazards such as the ultimate fate of the added chlorate and its impact on animal health if it remains in the silage.

Denitrification is a microbial process wherein NO_3_^-^ is reduced to N_2_ via intermediates including N_2_O. Factors such as a low C/N ratio have been reported to lead to N_2_O accumulation during denitrification (43, 44). The addition of acetate as an external carbon source, effectively increasing the C/N ratio, resulted in a slight reduction in N_2_O production (Fig. 2a and SI Table S1). This suggests that N_2_O production in conserved forage may be influenced, at least in part, by a low C/N ratio. Moreover, the findings also indicate that denitrification inhibitors, such as chlorate, can be combined with an external carbon source, such as acetate, as an effective additive to mitigate N_2_O emissions from the forage conservation process.

### N_2_O production and its relationship with nutritional parameters

N_2_O production was closely correlated with most parameters related to protein and amino acids, including NO_3_^-^-N (Fig. 3), presumably the main source of N_2_O production. The NO_3_^-^-N content in forages varies with the stage of plant maturity (45), and both alfalfa varieties harvested at later growth stages, which produced significantly less N_2_O (*p* < 0.05) (Fig. 1b), contained lower NO_3_^-^-N levels (SI Fig. S1). Reports have also demonstrated that nitrogen fertilization directly impact NO_3_^-^-N content (45), implying that nitrogen fertilization, especially preceding harvest, may contribute to higher N_2_O production at ensiling. Further studies are needed to investigate the effects of nitrogen fertilization schedule on N_2_O production. Simple changes in agricultural practice may reduce N_2_O emissions.

A source of organic carbon is an important component of denitrification, serving as an electron donor. Many studies have shown that external carbon sources, such as methanol, ethanol, and acetate, stimulate denitrification and usually reduce N_2_O production (46). Consistent with these findings, our study showed that the addition of acetate reduced N_2_O production (Fig. 2a). However, carbohydrate-related parameters, such as ADF, aNDF, lignin, starch, and ethanol-soluble carbohydrates, did not correlate with N_2_O production (Fig. 3), which could be due to the recalcitrance of these carbon sources. Similarly, denitrification was promoted by plant-based carbon substrates, such as rice straw, but there was a significant lag before denitrification became active (47).

### Environmental implications

What gets measured gets managed. The first step in reducing GHG emissions is to measure them. The US EPA publishes an annual report titled “Inventory of U.S. Greenhouse Gas Emissions and Sinks,” which estimates total GHG emissions by source across all sectors of the economy at the national level (1). Notably, agriculture is the largest contributor to N_2_O emissions in the US, accounting for 80% in 2021. The EPA monitors major sources in the agricultural sector, including agricultural land management, manure management, and the field burning of agricultural residues (Table 1) (1). In our study, 18.2 (*±*1.9), 169.7 (*±*10.9), and 4.8 (*±*0.2) g CO_2_ eq. per kg_DM-forage_ of N_2_O were produced from maize, alfalfa, and sorghum, respectively (Fig. 1a). According to the Crop Production 2022 Summary of the United States Department of Agriculture National Agricultural Statistics Service, the total production volumes of maize, alfalfa, and sorghum for silage in 2022 were 128.6, 17.4, and 5.6 MMT, respectively, comprising 93% of the total silage production combined (14). Assuming a similar amount of N_2_O can be produced from each crop, the total N_2_O emission potential amounts to 5.3 MMT CO_2_ eq.. This makes forage conservation the third largest N_2_O emitter in the agricultural sector, surpassing the field burning of agricultural residues by a factor of 30 (Table 1). Notably, N_2_O emissions from silage of uncategorized crops (total production volume: 10.7 MMT in 2022, comprising 7% of the total silage production) were not considered in this study (14). Again, the first step to reducing GHG emissions is to measure them, as policymakers and decision-makers use GHG inventories to develop strategies and track progress in GHG emission reduction efforts (48, 49).

**Table 1.** N_2_O emissions from the agriculture sector (MMT^a^ CO_2_ eq.) according to the Inventory of U.S. Greenhouse Gas Emissions and Sinks (1990–2021) by the EPA.

| Gas/Source | 2005 | 2019 | 2020 | 2021 | 2022 |
| --- | --- | --- | --- | --- | --- |
| Agricultural Soil Management | 291.5 | 309.3 | 290.5 | 294.0 | <sup>b</sup> |
| Manure Management | 14.5 | 17.4 | 17.5 | 17.4 | <sup>b</sup> |
| Field Burning of Agricultural Residues | 0.2 | 0.2 | 0.2 | 0.2 | <sup>b</sup> |
| **Forage Conservation<sup>c</sup>** | **5.6** | **5.6** | **5.7** | **5.8** | **5.3** |

Note: Totals may not sum due to independent rounding
<sup>a</sup> MMT: million metric tons
<sup>b</sup> data not available
<sup>c</sup> hypothetical estimation based on this study

### Limitations and Perspectives

There are, admittedly, limitations to our assessment, especially in extrapolating our data to the national scale. The microbial N_2_O production, like any other microbial processes, is sensitive to various environmental factors such as temperature and nutritional parameters, which could result in underestimation or overestimation of the outcomes. There are additional uncertainties associated with the heterogeneity of farmers’ practices, such as harvest time, inoculant use, moisture content (i.e., wilting), and oxygen exposure. Therefore, N_2_O emission measurements from actual silage fermentation systems spanning the range of environmental and management variation across the US are warranted to achieve more accurate estimation results. With that, this work aims to draw attention to the silage process as an overlooked but abundant source of N_2_O in the agricultural sector.

In this study, the use of chlorate was proposed as a simple and effective remedy to reduce N_2_O emissions significantly (Fig. 2a), and additional experiments demonstrated that lower concentrations as low as 0.01% (w/w) still achieved 92% N_2_O reduction (Fig. 2c). The chlorate (as sodium salt) price at the end of 2023 in the US was 795 USD per ton (50). The estimated cost to add 0.01% (w/w) chlorate as an additional silage additive is 0.08 USD per ton forage (dry weight), which is only approximately 5-8% compared to the silage inoculant cost (i.e., 1-1.5 USD per ton). Assessing the social cost of GHG emissions has become a common yardstick for estimating the benefits of formulating and implementing abatement policies (51). The social cost estimates of N_2_O shown in recent studies range from 16 to 174 USD per kg-N_2_O-N (52–54). Assuming that chlorate (0.01%, w/w) is added to achieve 90% N_2_O reduction (Fig. 2c), the social cost saving could be 81-882 million USD with the expected total cost of 12 million USD for applying chlorate as an additive.

## Materials and Methods

### Forage sample preparation

Maize (Dyna Gro D53VC55RIB) and sorghum (Dyna Gro Super Sile 20) were harvested from a private farm near Manhattan, KS (39°38’N, 96°88’W) at their optimal maturity stages (i.e., 2/3 milk line and soft dough stages, respectively). The plants were chopped to a theoretical length of 2 cm using a standard forage harvester without inoculation. Two distinct alfalfa varieties, HVX MegaTron (WinField United L.L.C., Arden Hills, MN, USA) and HybriForce 3400 (Dairyland Seed Co.), were grown in experimental plots at the Kansas State University Agronomy Research Farm in Manhattan, Kansas (39°20’s N, 96°59’W). Each variety was harvested at two different stages of maturity (mid-bud and early flowering) using a sickle bar mower. The harvested alfalfa was subsequently chopped to a length of 2 cm using a stationary forage chopper.

### Experimental design

We used a simulated silage model known as mini silos. These mini-silos comprised a 1-L glass jar (55) connected to a 3-L Tedlar bag (SI Fig. S4. Each crop was ensiled with four treatments: 1) no inoculant (control, I^-^), 2) crop-specific commercial silage inoculant (brand names omitted for confidentiality) following the manufacturer’s instructions (I^+^), 3) inoculant + chlorate (0.1%, w/w, potassium salt) (I^+^ Ch^+^), and 4) inoculant + acetate (0.1%, w/w, sodium salt) (I^+^ Ac^+^). Chlorate was added to inhibit denitrifiers (40, 41), and acetate was added to increase the initial C/N ratio (43, 44). Separate inoculant, chlorate, and acetate solutions were prepared and sprayed onto the plants. The initial moisture content was measured using the conventional microwave oven method (56) and adjusted to 70% (w/w, wet weight basis) by adding deionized water to the treatment solutions. All crops were packed in mini-silos with a bulk density of 650 kg/m^3^ (41 lb/ft3) (57), and each silo contained 650 g of the crops (wet weight), which was equivalent to 195 g_DM_. The mini-silos were incubated at 30°C in the dark, and N_2_O production was monitored regularly for up to 4 weeks. Gas bags were removed on Days 1, 2, 3, 5, and 20 to characterize temporal variations in the volume and composition of gas production and replaced with new bags at each sampling. Fresh feed samples were collected and stored at -80°C for chemical analysis. Additionally, four mini-silos were prepared for each treatment and sacrificed on Days 1, 3, 5, and 20 for molecular analysis. A subset of the samples (5 g) was preserved in 5 mL RNA preservation solution (RNAprotect, QIAGEN, Hilden, Germany) and stored at -80°C. Each treatment comprised three replicates of the mini-silos.

### Contribution of nitrification to N_2_O emissions

To investigate the relative contribution of nitrification to N_2_O production, mini-silos were prepared with alfalfa (HVX Megatron) harvested at the early flowering stage, and acetylene (10 Pa) (36) was added at the beginning of incubation. In addition, the mini-silos were intermittently exposed to oxygen on Days 3, 5, and 10, achieved by supplying 20 mL of air (4 mL of oxygen) through a stainless-steel tube that reached the center of the mini-silos (Supplementary Fig. 3). Oxygen was added to separate bottles on each injection date.

### Analytical methods

The total gas production volume was measured using the water displacement method. N_2_O was quantified using an Agilent 7890 gas chromatograph with an electron capture detector and an HP-PLOT/Q column (30 m × 0.53 mm × 40 m). Fresh forage samples were sent to Rock River Laboratory, Inc., Watertown, WI, for nutritional analysis. The following parameters were examined: crude protein (CP), total amino acid, NH3-N content, acid detergent fiber (ADF), amylase-treated neutral detergent fiber (aNDF), lignin, starch, and ethanol-soluble carbohydrate (ESC). NO_3_^-^-N content (% of DM) was measured at the Kansas State University Soil Testing Laboratory.

### Nucleic acid extraction and quantitative PCR

Microbial nucleic acid extraction from plant material, particularly from the epiphytic phyllosphere, poses challenges due to plant-derived biomolecules such as proteins and nucleic acids (58, 59). In this study, we used a microbial DNA extraction method optimized for silage samples, as detailed in our previous study (60). Briefly, for DNA extraction, five grams of the sample was suspended in 45 mL of sterile 0.85% NaCl solution, shaken on a rotary shaker at 120 rpm for 2 h at room temperature, filtered through two layers of gauze cloth to remove large plant debris, and centrifuged at 12,000 g for 15 min at 4°C (61). The supernatant was discarded, and the pellet was stored at -80°C until subsequent DNA extraction. The RNA preservation solution containing the sample was shaken on a rotary shaker at 120 rpm for 2 h at room temperature. Four milliliters of the supernatant were collected, pelleted by centrifugation at 12,000 g for 15 min at 4°C, and immediately subjected to RNA extraction. DNA and RNA extractions from the pellets were performed using the DNeasy and RNeasy PowerSoil kits (Qiagen, Hilden, Germany), following the manufacturer’s instructions, with slight modifications. Cells were lysed by bead beating at 20°C for 5 min. RNA samples were subjected to DNase treatment (ezDNase™, Invitrogen, CA, USA) and reverse transcribed into complementary DNA (cDNA) using SuperScriptIV Reverse Transcriptase (Thermo Fisher Scientific, Waltham, MA, USA). PCR was conducted with universal 16S rRNA gene primers on DNase-treated RNA samples to confirm the absence of DNA contamination. Target genes and transcripts were quantified by qPCR using a CFX Opus 96 Real-Time PCR System (Bio-Rad, Hercules, CA, USA) in 20 μL reaction mixtures containing 10 μL of SsoAdvanced™ Universal SYBR Green Supermix (Bio-Rad, Hercules, CA, USA), 300 nM of each primer, and 2 μL of template DNA. Quantification was performed using standard curves prepared from serial 10-fold dilutions of cloned plasmids or double-stranded synthetic DNA fragments (gBlocks®, Integrated DNA Technologies, Coralville, IA, USA) in triplicate. Detailed information on primer sequences, standard sequences, and detection limits can be found in SI Table S2.

### Statistical analysis

Statistical analyses were conducted using the MIXED procedures of SAS/STAT software, Version 9.4 (SAS Institute Inc., Cary, NC, USA). A two-way ANOVA was applied to the N_2_O emission data, crop, treatment, and their interaction term as fixed effects. A Bonferroni multiplier adjustment was performed when comparing N_2_O emissions between crops in each treatment group or treatments in each crop group (conditional pairwise comparison). Probability values of *p* < 0.05 (2-tailed) were considered statistically significant for all comparisons. The correlation coefficients between N_2_O production and nutrient variables were calculated as Pearson correlation coefficients. A Pearson correlation coefficient > 0.8 or < -0.8 with *p* < 0.05 (two-tailed) was deemed indicative of a strong correlation between the two variables.

## Data Availability

All data are included in the article and Supplementary material.

## ACKNOWLEDGMENTS

Please include your acknowledgments here, set in a single paragraph. Please do not include any acknowledgments in the Supporting Information, or anywhere else in the manuscript. Support for this research was provided by the Kansas NSF EPSCoR (OIA-1656006) and NSF CAREER Program (2144189).

