## Supplementary material for "Forage conservation is a neglected nitrous oxide source"

### This PDF file includes:

Figs. S1 to S4
Tables S1 to S2
SI References

**Table S1. Least square means for $N_2O$ emissions (g-$CO_2$ eq. per $kg_{DM\text{-}forage}$) with standard error (SE) of the two-way analysis of variance (ANOVA) model under variance heterogeneity.†, ‡, #**

| | | Treatments | | |
|---|---|---|---|---|
| Crops | $I^-$ | $I^+$ | $I^+ Ch^+$ | $I^+ Ac^+$ |
| $Cv_1 Hv_1$ | $144.14 \pm 10.90^a$ | $130.33 \pm 10.90^a$ | $8.90 \pm 10.90^b$ | $114.07 \pm 10.90^a$ |
| $Cv_2 Hv_1$ | $169.74 \pm 10.90^a$ | $177.72 \pm 10.90^a$ | $9.01 \pm 10.90^c$ | $93.84 \pm 10.90^b$ |
| $Cv_1 Hv_2$ | $78.99 \pm 7.77^a$ | $91.82 \pm 7.77^a$ | $4.15 \pm 7.77^b$ | $71.23 \pm 7.77^a$ |
| $Cv_2 Hv_2$ | $83.91 \pm 7.77^a$ | $88.79 \pm 7.77^a$ | $0.83 \pm 7.77^b$ | $78.71 \pm 7.77^a$ |
| Maize | $18.24 \pm 1.87^a$ | $20.57 \pm 1.87^a$ | $2.70 \pm 1.87^b$ | $16.96 \pm 1.87^a$ |
| Sorghum | $4.79 \pm 0.22^a$ | $2.46 \pm 0.22^b$ | $0.07 \pm 0.22^c$ | $0.52 \pm 0.22^c$ |

[a, b, c]Means in the same row with different superscript letters differ significantly ($P < 0.05$).

[†]Treatments are no inoculant ($I^-$), crop-specific commercial silage inoculant ($I^+$), inoculant and chlorate ($I^+ Ch^+$), and inoculant and acetate ($I^+ Ac^+$). The notations $Cv_1$ and $Cv_2$ indicate two alfalfa cultivars, HVX MegaTron and HybriForce 3400, respectively. The notations $Hv_1$ and $Hv_2$ indicate samples harvested at two different maturity stages, i.e., mid-bud and early-flowering, respectively.

[‡]Results are reported as lease square means ± standard error (SE) for each combination group of crop and treatment.

[#]P values of two-way analysis of variance (ANOVA) under variance heterogeneity for crop, treatment, and crop by treatment interaction are < 0.01, <0.01, and <0.01, respectively.

**Table S2. Quantitative PCR primers, standards, and detection limits of each assay.**

| Target gene | Primer set | Primer sequences (5'-3') | Quantification range |
|---|---|---|---|
| 16S rRNA [1] | 8F<br>1492R | AGAGTTTGATCCTGGCTCAG<br>TACCTTGTTACGACTT | $10^2$-$10^9$ |
| *napA* [2] | V67m<br>V17m | AAYATGGCVGARATGCACCC<br>GRTTRAARCCCATSGTCCA | $10^2$-$10^9$ |
| *narG* [3] | narG-F<br>narG-R | TCGCCSATYCCGGCSATGTC<br>GAGTTGTACCAGTCRGCSGAYTCSG | $10^2$-$10^9$ |
| *nirK* [4] | nirKC1-F<br>nirKC1-R | ATGGCGCCATCATGGTNYTNCC<br>TCGAAGGCCTCGATNARRTTRTG | $10^4$-$10^9$ |
| *nirS* [5] | nirS-1F<br>nirS-6R | CCTAYTGGCCGCCRCART<br>CGTTGAACTTRCCGGT | $10^3$-$10^9$ |
| *qnorB* [6] | qnorB-2F<br>qnorB-5R | GGNCAYCARGGNTAYGA<br>ACCCANAGRTGNACNACCCACCA | $10^2$-$10^9$ |
| Clade I nosZ [7] | nosZ-1F<br>nosZ-1R | WCSYTGTTCMTCGACAGCCAG<br>ATGTCGATCARCTGVKCRTTYTC | $10^2$-$10^9$ |
| Clade II *nosZ* [8] | nosZ 1374F<br>nosZ 1577R | CCBYTBCAYACSCARTTYG<br>TGSGASAGCTTGTTSAGSS | $10^2$-$10^9$ |
| Bacterial - *amoA* [9] | amoA-1F<br>amoA-2R | GGGGTTTCTACTGGTGGT<br>CCCCTCKGSAAAGCCTTCTTC | $10^2$-$10^9$ |
| Archaeal - *amoA* [10] | arch-amoAF<br>arch-amoAR | STAATGGTCTGGCTTAGACG<br>GCGGCCATCCATCTGTATGT | $10^2$-$10^9$ |

Abbreviations for degenerate nucleotide positions are as follows: R =A or G; K = G or T; M =A or C; S =C or G;W=A or T; Y =C or T; B =C, G, or T; D = A, G, or T; V =A, C, or G; H =A, C, or T. References for the primers and probes are included in the first column.

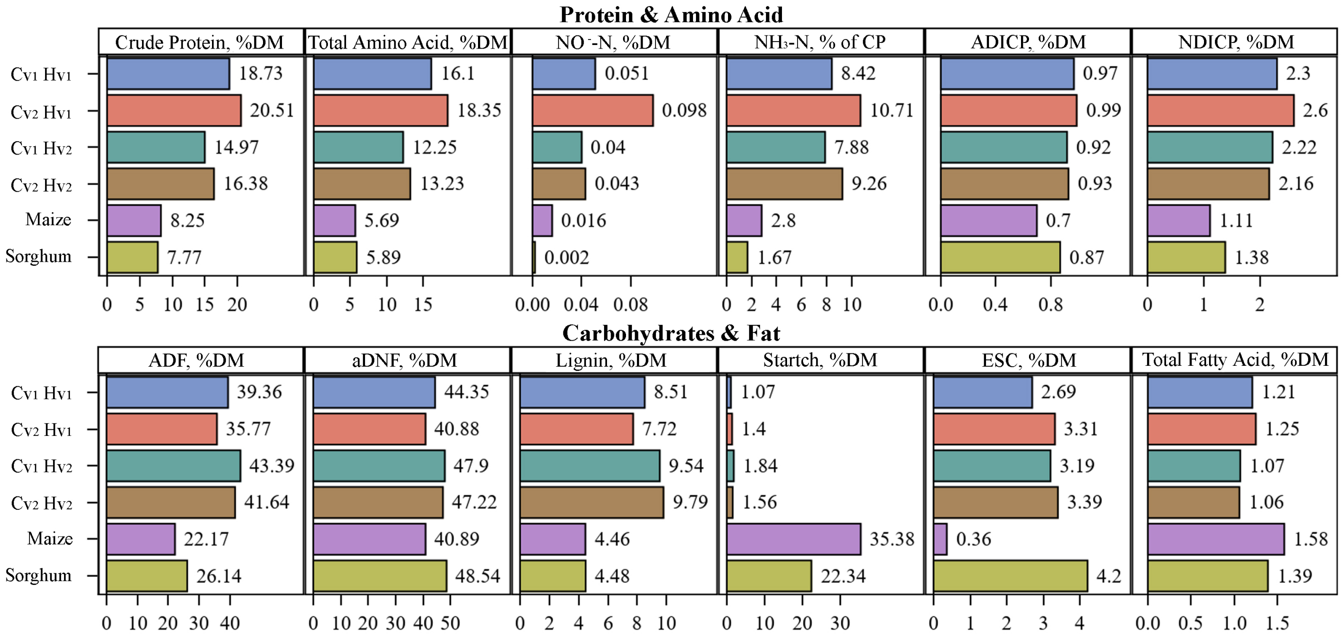


**Fig. S1.** Nutritional parameters of fresh maize, alfalfa (two distinct varieties harvested at two different stages of maturity), and sorghum samples. The labels $Cv_1$ and $Cv_2$ denote two alfalfa varieties, HVX MegaTron and HybriForce 3400, respectively. The labels $Hv_1$ and $Hv_2$ denote alfalfa samples harvested at the mid-bud and early flowering stages, respectively. DM: dry matter; CP: crude protein; ADICP: acid detergent-insoluble CP; NDICP: neutral detergent-insoluble CP; ADF: acid detergent-treated fiber; aNDF: amylase-treated neutral detergent fiber; ESC: ethanol-soluble carbohydrates. The numerical annotations next to the individual bars are their respective measured values.

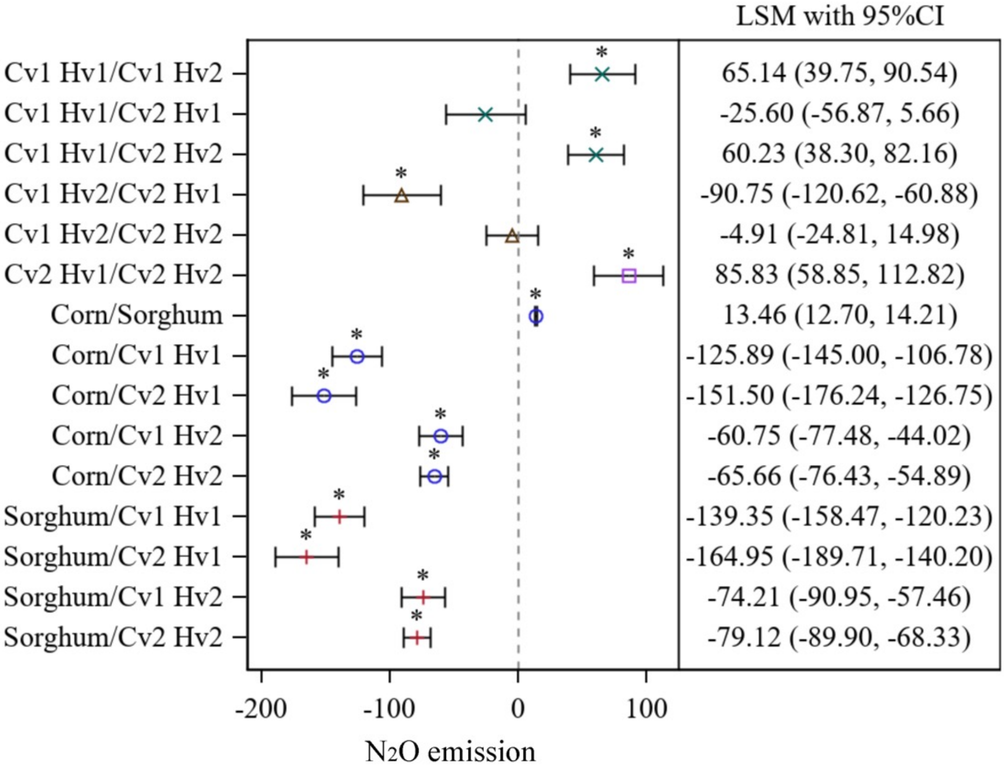


**Fig. S2.** Pairwise comparisons $N_2O$ emissions for crops under the controls (I⁻) group. Results are reported as lease square means for $N_2O$ emissions with 95% confidence intervals. '*' indicates a significant difference observed between the two crop groups.

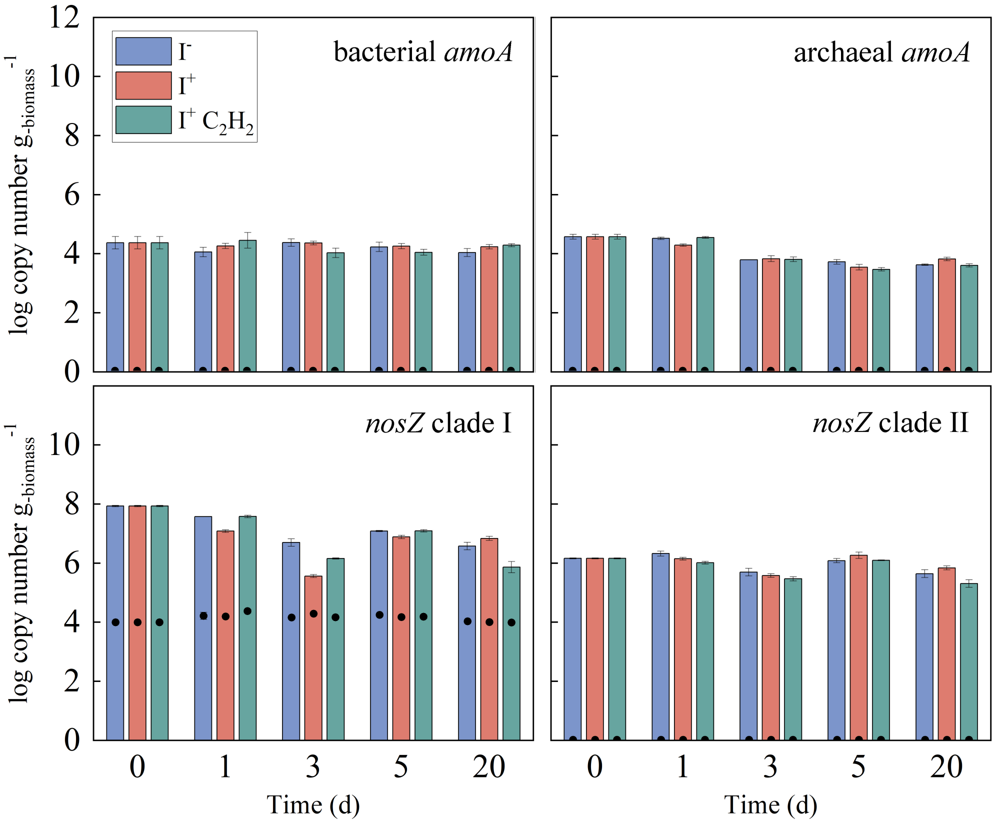


**Fig. S3.** Effect of acetylene ($C_2H_2$) on the abundance of bacterial/archaeal *amoA* genes and clade I/II *nosZ* genes (bar) and their transcripts (black circle). The error bars represent the standard deviations of the triplicate incubations. Error bars may not be visible if their magnitude is smaller than the symbols.

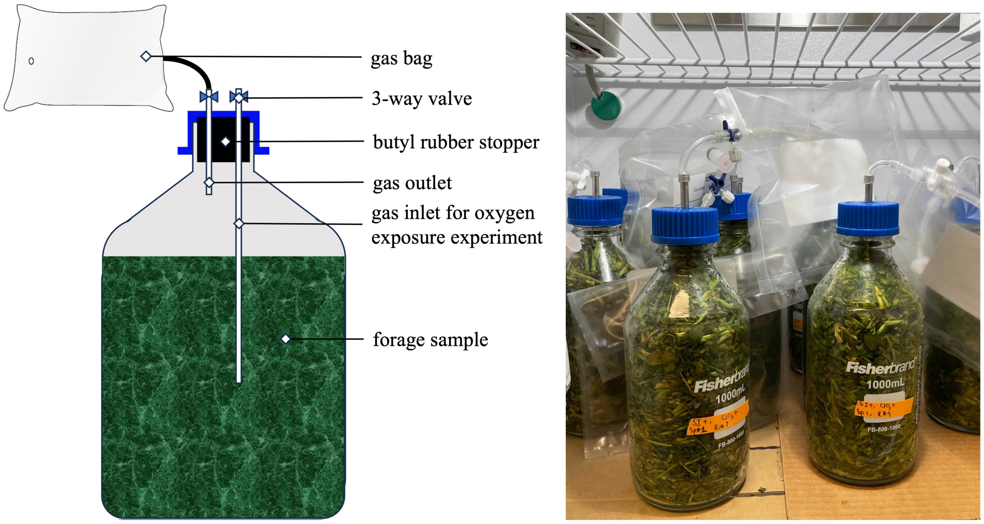


**Fig. S4.** A schematic (left) and photo (right) of mini-silos used in this study. The gas inlet was only used for the oxygen exposure experiment.
